# Genome mining reveals a sporulation-associated protein with ferredoxin–NADP^+^ reductase activity in *Clostridium pasteurianum*: structural and kinetic characterization

**DOI:** 10.64898/2026.08.07.743380

**Authors:** Weigao Wang, Qianqiao Liu, James R. Swartz

**Author notes:** Correspondence: J. R. Swartz.

## Abstract

Ferredoxin–NADP^+^ reductases (FNRs) are ubiquitous flavoenzymes that catalyse the reversible transfer of electrons between iron–sulfur ferredoxins and the pyridine nucleotide pool, thereby occupying a central position in diverse redox metabolic pathways including photosynthesis, nitrogen fixation, and detoxification of reactive oxygen species. Although FNR activity was demonstrated in cell extracts of *Clostridium pasteurianum* more than five decades ago, the gene encoding this activity has remained unidentified. In the present study, a systematic bioinformatic screen of all 3,797 predicted proteins from the *C. pasteurianum* genome was conducted using conserved FAD- and NAD(P)^+^-interacting residues from structurally characterised reductases as search templates. This analysis identified a single candidate, AQ984_05830, which is annotated as a sporulation protein but possesses all six predicted cofactor-interacting residues. Heterologous expression and cytochrome *c* reduction assays confirmed ferredoxin-dependent reductase activity, with a wild-type *k*_cat_ of 0.007 min^−1^—a value orders of magnitude lower than those reported for canonical FNRs. A parallel genome-wide screen further revealed a repertoire of ferredoxin-like carriers, suggesting that *C. pasteurianum* distributes hydrogen-derived electrons among multiple ferredoxins to serve diverse metabolic fates, of which NADP reduction by CpFNR is one. Alanine scanning mutagenesis of five predicted cofactor-interacting residues revealed that K68A and K73A mutations abolished activity, whereas T64A, T185A and S202A mutations improved catalytic efficiency (*k*_cat_/*K*_m_) for NADH by 14 to 18 folds. AlphaFold structure prediction combined with SwissDock and ClusPro molecular docking simulations placed the FAD binding site centrally between the NAD(P)H and ferredoxin binding domains, consistent with the expected electron relay architecture. Structural analysis of the beneficial mutations suggests that disruption of hydrogen bonds flanking a flexible coil (residues 186–199) propagates conformational effects to the NAD(P)H binding loops, rationalising the improved substrate affinities. These findings expand the known functional diversity of the FNR superfamily and suggest an unrecognised role for redox regulation during endospore formation in *C. pasteurianum*.

## Introduction

Ferredoxin–NADP^+^ reductases (FNRs, EC 1.18.1.2) constitute a widely distributed family of FAD-containing oxidoreductases that catalyse the reversible transfer of electrons between low-potential iron–sulfur ferredoxins and the pyridine nucleotide pool [1,2]. In photosynthetic organisms, plastidic FNRs supply the NADPH consumed by the Calvin cycle, whereas in bacteria and archaea, non-plastidic FNRs participate in nitrogen fixation, hydrogen production, steroid hydroxylation and the detoxification of reactive oxygen species [2,3]. The breadth of these functions underscores the evolutionary importance of the FNR superfamily and has motivated extensive structural and mechanistic studies over the past four decades.

From a structural perspective, FNRs share a conserved two-domain architecture comprising an FAD-binding domain and an NAD(P)^+^-binding domain [4–6]. The FAD cofactor is deeply buried within the protein, its isoalloxazine ring stabilised by hydrogen bonds and van der Waals contacts with conserved residues in the binding pocket, while the NAD(P)^+^-binding domain engages the pyridine nucleotide through specific interactions involving conserved lysine, arginine, serine and threonine residues that position the nicotinamide ring for hydride transfer to the flavin [5,7,8]. It has been demonstrated that mutations at these conserved positions frequently produce dramatic changes in catalytic properties, as shown by extensive mutagenesis studies on the *Anabaena* PCC 7119 FNR and related enzymes [7,9–11]. The electron transfer interface between FNR and its ferredoxin partner is governed by electrostatic complementarity between acidic residues on ferredoxin and basic residues on FNR, supplemented by hydrophobic contacts that optimise the orientation for efficient electron transfer [6,7,21,23,24].

*Clostridium pasteurianum* is a nitrogen-fixing obligate anaerobe that has served as a model organism for studies of ferredoxin biochemistry since the pioneering work of Mortenson in the 1960s [12]. In a seminal study, Jungermann and co-workers demonstrated ferredoxin–NADP^+^ reductase activity in cell extracts of *C. pasteurianum* [12]; however, despite more than five decades of research on the redox enzymes of this organism, the gene encoding this activity has never been identified. This gap has limited our understanding of the electron transfer network in *C. pasteurianum* and has precluded the rational engineering of complete cofactor regeneration cascades from this industrially relevant organism.

In recent years, it has become increasingly apparent that many proteins possess secondary or ‘moonlighting’ functions that are unrelated to their annotated primary role [32]. Such moonlighting activities are particularly difficult to identify by sequence homology alone, as the secondary function may be mediated by structural features that are not reflected in the primary sequence annotation. Sporulation in clostridia is a complex developmental programme triggered by environmental stress [35], and optimal redox conditions have been proposed to promote the physiological processes necessary for endospore formation [36]. Whether dedicated redox enzymes participate in regulating the intracellular redox environment during sporulation has remained an open question.

In the present study, we describe the identification and biochemical characterisation of a novel FNR from the *C. pasteurianum* genome. Using a systematic bioinformatic screen based on conserved cofactor-interacting residues from structurally characterised reductases, we identified a single candidate—a protein annotated as a sporulation factor—that possesses all predicted FAD and NAD(P)^+^ contact residues. We confirmed its FNR activity biochemically, mapped the contributions of individual cofactor-interacting residues through alanine scanning mutagenesis, and employed AlphaFold-predicted structures combined with molecular docking to rationalise the observed kinetic effects. Our findings reveal an unexpected functional link between sporulation and redox regulation in *C. pasteurianum* and expand the known diversity of the FNR superfamily.

## Materials and methods

### Chemicals and reagents

All buffer chemicals were obtained from Sigma-Aldrich (St Louis, MO, USA) unless stated otherwise. PCR components, T4 polymerase and Gibson Assembly mix were purchased from New England Biolabs (Ipswich, MA, USA). Synthetic gene fragments (gBlocks) and oligonucleotides were obtained from IDT DNA (Coralville, IA, USA). Plasmid DNA was isolated using the Plasmid Plus Midi Kit (Qiagen, Hilden, Germany), and PCR products were purified with the DNA Clean & Concentrate kit (Qiagen). Strep-Tactin XT Sepharose chromatography resin was purchased from Cytiva Life Sciences (Marlborough, MA, USA).

### Molecular cloning and site-directed mutagenesis

DH10β cells (NEB) were used for cloning and plasmid propagation in LB medium supplemented with 100 μg·mL^−1^ ampicillin. BL21(DE3) and BL21(DE3)(Δ*ISCR*) cells were used for expression of the ferredoxin NADP^+^ reductase and ferredoxin, respectively. The pET21b vector was ligated with the gene of interest using Gibson Assembly. Alanine substitutions (T64A, K68A, K73A, T185A, S202A) were introduced by site-directed mutagenesis. Primers and gBlock sequences are summarised in Table S1. All plasmids were verified by Sanger sequencing.

### Protein expression and purification

Expression and purification of CpI hydrogenase and CpFd were conducted as previously described [13]. BL21(DE3) cells harbouring the CpFNR gene were grown in 1 L of LB medium containing 100 μg·mL^−1^ ampicillin at 37 °C. At an OD_600_ of 0.4–0.6, IPTG was added to a final concentration of 0.5 mM, and incubation was continued at 28 °C for 16 h. Cells were harvested by centrifugation at 5000 *g* for 20 min and resuspended in a binding buffer (0.1 M Tris-HCl, 0.15 M NaCl, pH 8.0). Cell lysis was achieved by two passes through a homogeniser, followed by centrifugation at 10 000 *g* for 20 min. The supernatant containing C-terminally Strep-tagged CpFNR was applied to a 10 mL Strep-Tactin resin column, washed with five column volumes of binding buffer and eluted with elution buffer (0.1 M Tris-HCl, 0.15 M NaCl, 50 mM biotin, pH 8.0). Purity was assessed by SDS-PAGE.

### Enzyme kinetics

Enzyme kinetics were measured using a cytochrome *c* reduction assay as previously described [13]. All reaction components were prepared in an anaerobic glove box (COY Inc., Grass Lake, MI, USA). Reactions were conducted in UV-transparent half-area 96-well plates at room temperature. Horse heart cytochrome *c* (50 μM) served as the terminal electron acceptor, and NADH or NADPH served as electron donors. Reduction of cytochrome *c* was monitored at 550 nm in 100 mM Tris-HCl (pH 7.0). All experiments were performed in triplicate.

### Bioinformatic screening and sequence analysis

A total of 3798 amino acid sequences from the *C. pasteurianum* genome were extracted from NCBI using GeneBank Trans Extractor and aligned with structurally characterised FNRs—FdR from *Thermobifida fusca* (PDB: 6TUK) and BphA4 from *Pseudomonas* sp. (PDB: 2GQW)—using Clustal Omega. Conserved cofactor-interacting residues were identified, and candidate proteins were selected based on the presence of exact or chemically equivalent residues at all six key positions. In parallel, the same 3798 protein sequences were screened for ferredoxin-like electron carriers by identifying the bacterial-type [4Fe-4S] dicluster signature (CxxCxxCxxxCP), using the *C. pasteurianum* ferredoxin (CpFd; GenBank AOZ73678.1) as the reference sequence; motif-containing proteins were cross-referenced against their NCBI annotations and ranked by sequence length.

### Structure prediction and molecular docking

The tertiary structure of CpFNR was predicted using AlphaFold (v.2.3.2). Docking predictions for CpFNR with NADPH, NADH and FAD were performed using SwissDock [14,15], and the CpFd–CpFNR interaction was modelled using ClusPro [16]. Structural analyses were visualised using PyMOL [17].

## Results

### Genome-wide screening identifies a sporulation protein as the sole FNR candidate

Structural analysis of the template reductases FdR (6TUK) and BphA4 (2GQW) established six key cofactor-interacting positions: three residues that stabilise the FAD cofactor through hydrogen bonds with the adenosine ribose (E35/E41), ionic interactions with the proximal phosphate (R42/R48) and hydrogen bonds with the isoalloxazine ring (K47/K53); and three residues that stabilise the NAD moiety through hydrogen bonds with the ribose (E175/E186), hydrogen bonds with the ribose hydroxyl (S182/S195) and ionic interactions with the phosphate (R183). For each position, a set of permissible replacement residues was defined based on side-chain chemistry (Fig. 1).

**Figure 1.**
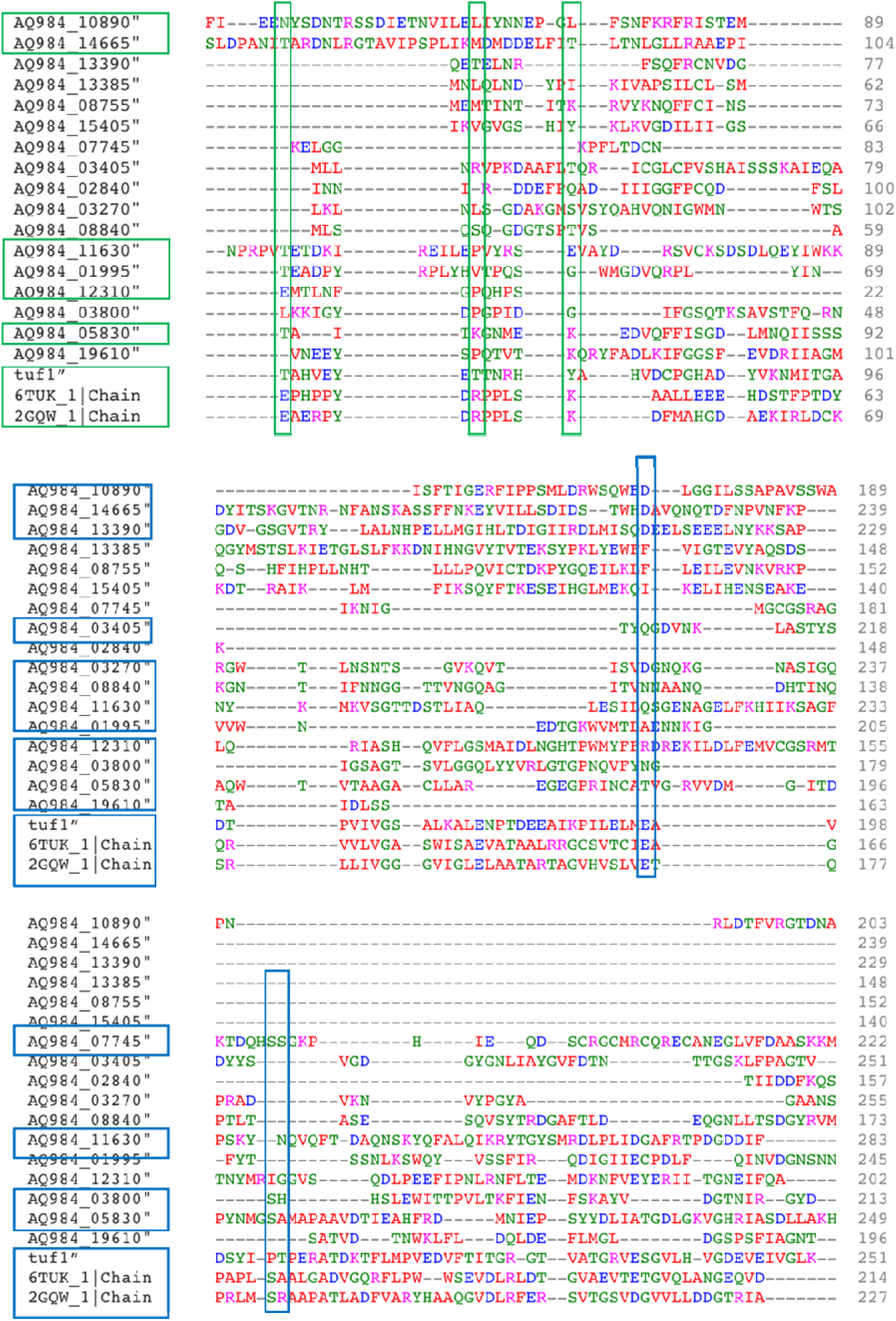
Sequence alignments between the template NAD reductase enzymes (6TUK and 2GQW) and selected protein sequences from the *Clostridium pasteurianum* genome. Residues with the potential for forming hydrogen bond interactions are highlighted. The FAD cofactor is primarily stabilized by glutamic acid, arginine, and lysine residues, while the NAD^+^ cofactor is mainly stabilized by glutamic acid, serine, and arginine residues. Proteins showcasing similar key residue features are also highlighted. Green boxes indicate FAD-binding residues, while blue boxes designate NAD-binding residues.

Screening of all 3798 *C. pasteurianum* proteins revealed that the majority possessed only one or two matching residues (Fig. 2). A single protein, AQ984_05830, was found to contain all six predicted cofactor-interacting residues at the corresponding alignment positions. It is noteworthy that this protein is annotated as a sporulation protein rather than a reductase. This protein was designated CpFNR and subjected to biochemical characterisation.

**Figure 2.**
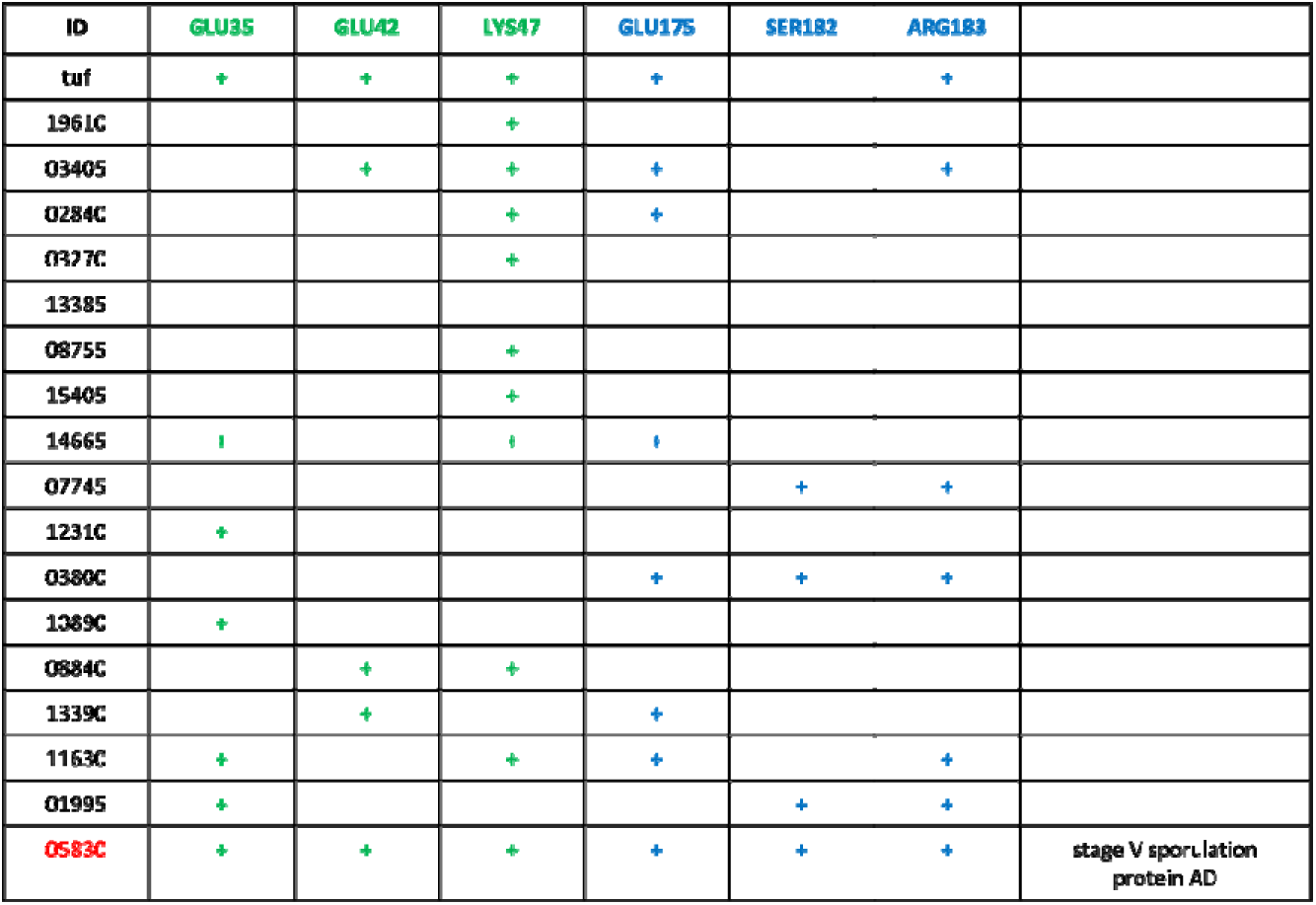
Summary of selected proteins in *C. pasteurianum* genome. Residues potentially forming interactions with FAD group were colored in green, and NAD group colored in blue.

To place CpFNR within the broader electron-transfer machinery of *C. pasteurianum*, we also surveyed the genome for ferredoxin-like proteins, the physiological electron donors of FNRs (Fig. 3). Of the 3798 proteins, 36 contained the bacterial-type [4Fe-4S] dicluster signature (CxxCxxCxxxCP), of which 16 were annotated as ferredoxins and 6 corresponded to bona-fide low-molecular-weight ferredoxins (≤200 aa). CpFd (AOZ73678.1) was recovered as the smallest of these carriers (56 aa) and served as the reference for the screen. The recovery of multiple ferredoxin-like carriers—16 annotated ferredoxins, six of them bona-fide low-molecular-weight proteins—alongside a single FNR candidate indicates that *C. pasteurianum* does not rely on one dedicated electron carrier. Instead, it appears to maintain a repertoire of ferredoxins capable of accepting the low-potential electrons released by [FeFe]-hydrogenase during H oxidation and partitioning them among distinct downstream acceptors. Within this network, CpFNR represents the branch that diverts a fraction of these electrons to the NADP pool, a role complementary to the better-characterised ferredoxin-dependent pathways of nitrogen fixation and hydrogen cycling in this organism.

**Figure 3.**
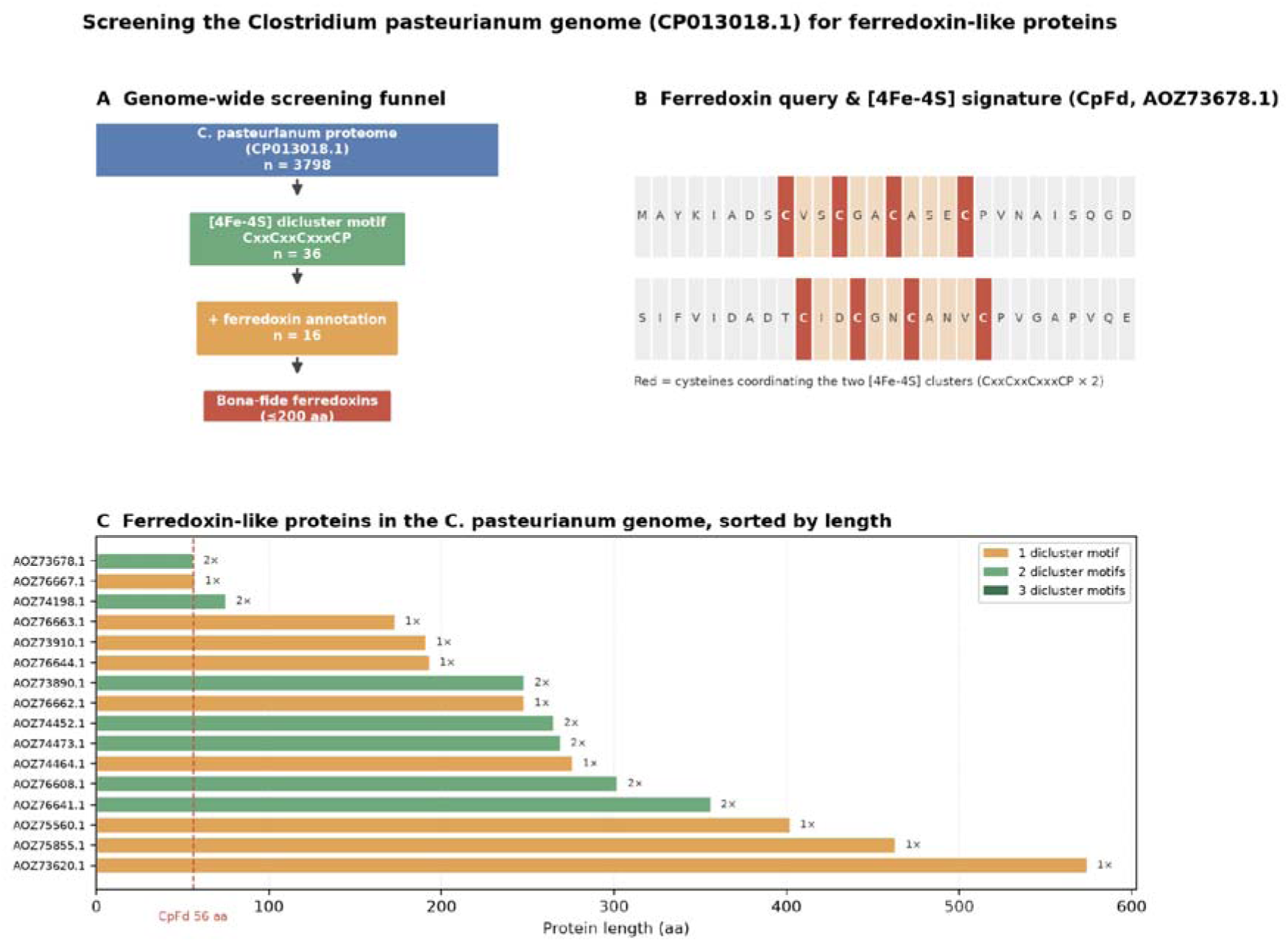
Genome-wide screening of the *Clostridium pasteurianum* genome (GenBank CP013019.1) for ferredoxin-like proteins. (A) Screening funnel showing the stepwise narrowing of candidates: the complete proteome (3798 proteins) was screened for the bacterial-type [4Fe-4S] dicluster signature (CxxCxxCxxxCP), yielding 36 motif-containing proteins, 16 of which are additionally annotated as ferredoxins, of which 6 are bona-fide low-molecular-weight ferredoxins (≤200 aa). (B) The C. *pasteurianum* ferredoxin query (CpFd, GenBank AOZ73678.1; 56 aa) shown residue-by-residue; cysteine residues coordinating the two [4Fe-4S] clusters are highlighted in red, and the two CxxCxxCxxxCP motifs define the ferredoxin signature used for the screen. (C) The 16 ferredoxin-annotated, motif-containing proteins sorted by length and coloured by the number of dicluster motifs; the dashed line marks the length of CpFd (56 aa) and the value beside each bar indicates the dicluster motif count.

### CpFNR exhibits confirmed but low FNR activity

To test whether AQ984_05830 possesses FNR activity, the protein was expressed heterologously, purified and assayed for its ability to catalyse ferredoxin-dependent cytochrome *c* reduction. As shown in Fig. 4, the wild-type enzyme exhibited a turnover number of 0.007 min^−1^, confirming FNR activity but at a rate that is orders of magnitude lower than those reported for canonical FNRs, which typically display *k*_cat_ values in the range of 49–349 s^−1^ [3].

**Figure 4.**
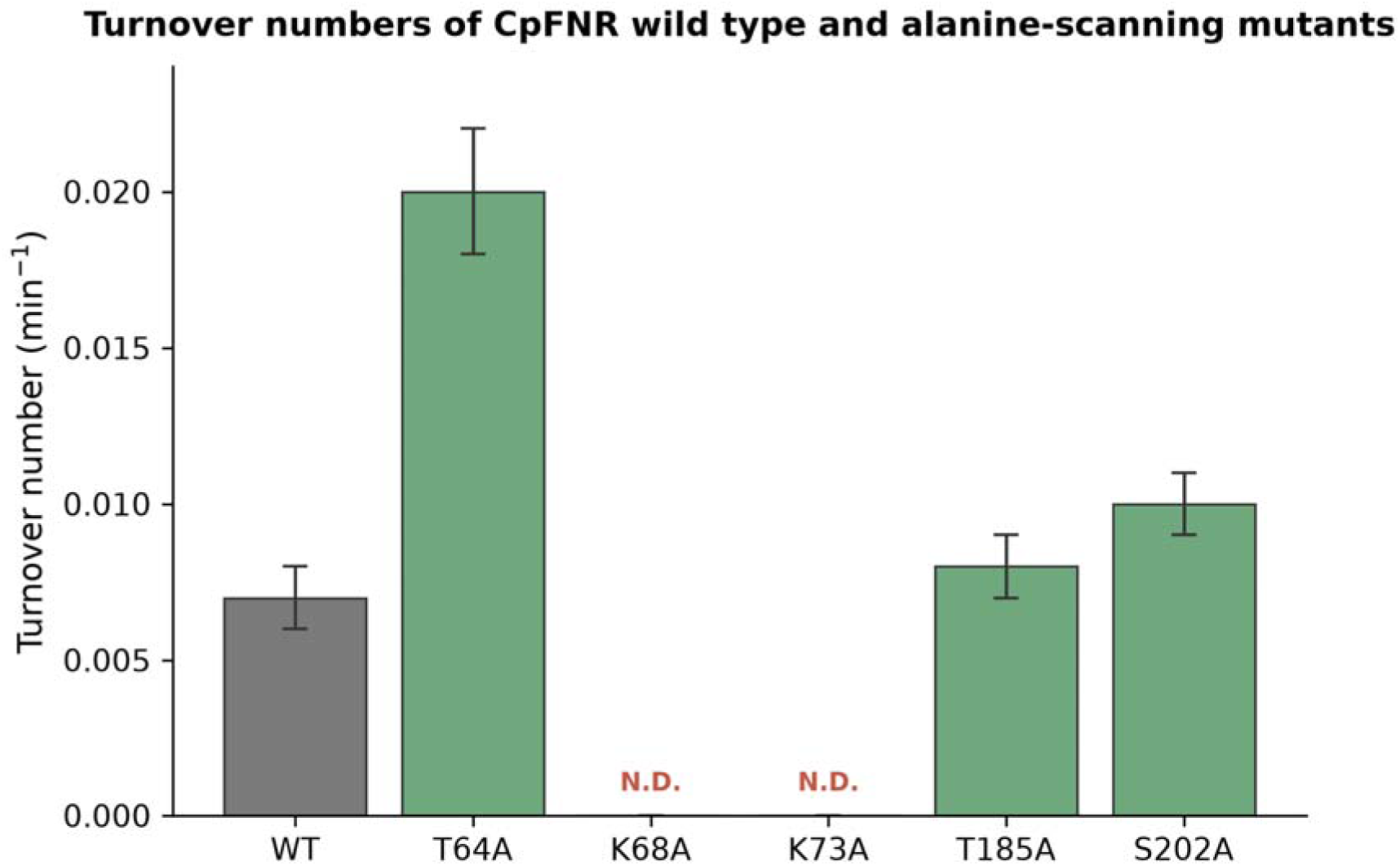
Turnover numbers of CpFNR wild type and alanine scanning mutants, measured by the rate of cytochrome *c* reduction. Error bars represent standard deviations of three independent replicates.

Alanine scanning mutagenesis was performed on five residues predicted to interact with FAD or NAD(P)H (Fig. 4). The K68A and K73A mutations completely abolished enzymatic activity, confirming the indispensable role of these lysine residues in catalysis. In contrast, T64A improved the turnover number approximately 3-fold to 0.02 min^−1^, T185A maintained wild-type levels and S202A produced a modest increase. The observation that mutations at predicted cofactor-interacting residues can enhance rather than diminish activity suggests that the hypothesised residue–cofactor contacts derived from sequence alignment may not accurately reflect the actual binding geometry of CpFNR.

### Differential effects of mutations on substrate specificity and catalytic efficiency

Steady-state kinetic parameters were determined for NADH, NADPH and CpFd with each active mutant (Table 1). Wild-type CpFNR exhibited a higher *k*_cat_ for NADH than for NADPH; however, the *K*_m_ for NADH was 7-fold higher than that for NADPH (428.5 vs. 64.87 μM), yielding a 4-fold higher catalytic efficiency (*k*_cat_/*K*_m_) for NADPH. This preference for NADPH over NADH is consistent with the cofactor specificity observed in other characterised FNRs [3,11,27].

**Table 1.** Kinetic parameters of CpFNR wild type and mutants.

| Substrate | | WT | T64A | T185A | S202A | $k_{cat}(min^{-1})/K_m(\mu M)$ | | | |
| --- | --- | --- | --- | --- | --- | --- | --- | --- | --- |
|  |  |  |  |  |  | WT | T64A | T185A | S202A |
| NADPH | $k_{cat}(min^{-1})$ | $0.123 \pm 0.006$ | $0.066 \pm 3 \times 10^{-4}$ | $0.066 \pm 0.003$ | $0.091 \pm 0.004$ | $0.002 \pm 9 \times 10^{-5}$ | $0.0041 \pm 2 \times 10^{-4}$ | $0.002 \pm 9 \times 10^{-5}$ | $0.002 \pm 1 \times 10^{-4}$ |
| | $K_m(\mu M)$ | $64.87 \pm 0.32$ | $16.3 \pm 0.08$ | $33.17 \pm 1.65$ | $41.63 \pm 2.08$ | | | | |
| NADH | $k_{cat}(min^{-1})$ | $0.211 \pm 0.001$ | $0.136 \pm 3 \times 10^{-4}$ | $0.27 \pm 0.01$ | $0.247 \pm 0.012$ | $0.0005 \pm 2 \times 10^{-5}$ | $0.0017 \pm 8 \times 10^{-5}$ | $0.009 \pm 4 \times 10^{-4}$ | $0.007 \pm 3 \times 10^{-4}$ |
| | $K_m(\mu M)$ | $428.5 \pm 2.1$ | $80.38 \pm 0.4$ | $29.78 \pm 1.48$ | $32.21 \pm 1.61$ | | | | |
| CpFdI | $k_{cat}(min^{-1})$ | $0.192 \pm 0.001$ | $9.243 \pm 0.046$ | $0.678 \pm 0.034$ | $0.59 \pm 0.03$ | $0.0133 \pm 6 \times 10^{-4}$ | $0.0288 \pm 0.001$ | $0.157 \pm 0.008$ | $0.07 \pm 0.003$ |
| | $K_m(\mu M)$ | $14.5 \pm 0.07$ | $321.5 \pm 1.6$ | $4.31 \pm 0.21$ | $8.27 \pm 0.41$ | | | | |

The T64A mutation decreased *k*_cat_ values for both pyridine nucleotides but improved *k*_cat_ for CpFd by 9-fold. Notably, T64A substantially improved affinity for NADPH and NADH while decreasing affinity for CpFd by more than 20-fold. Both T185A and S202A significantly improved affinity for all three substrates, with catalytic efficiency for NADH increasing more than 7-fold for both mutants. T64A was the only mutant that improved catalytic efficiency for NADPH, while T185A and S202A maintained wild-type levels for this substrate. These differential effects suggest that the mutations perturb distinct structural elements within the enzyme.

### Molecular docking reveals an unusual overlap between cofactor binding sites

To elucidate the structural basis of CpFNR catalysis, molecular docking simulations were performed using the AlphaFold-predicted structure. The docking results revealed that NADH and NADPH occupy partially overlapping but distinct binding sites on CpFNR (Fig. 5). The FAD binding site was found to overlap with the NAD(P)H site on the second loop (residues 309–313), a feature that contrasts with the spatially separated binding domains observed in canonical FNRs such as FdR9, PuR and BphA4 [4,5,18–20]. The second loop displays a characteristic **<u>G</u>**ALL**<u>S</u>**PL**<u>S</u>** motif, which differs from the GXSXXS motif identified in FdR9 [4], suggesting evolutionary divergence in cofactor recognition.

**Figure 5.**
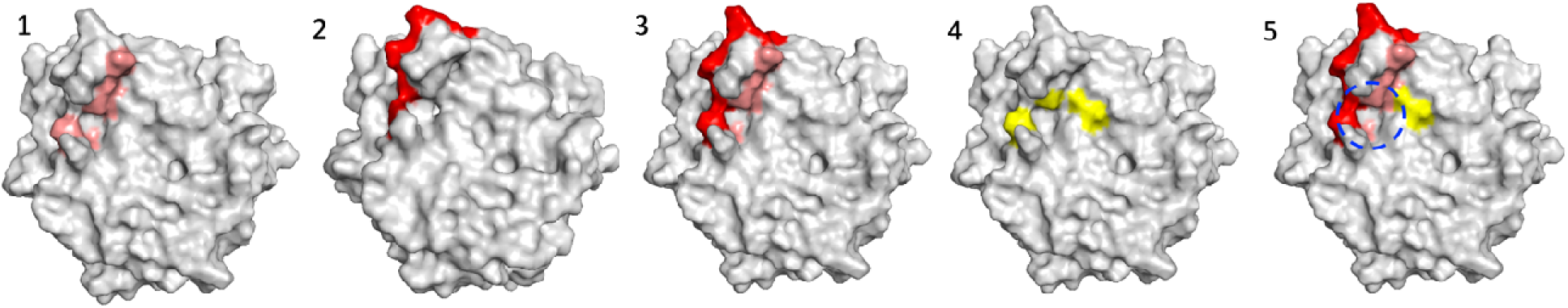
Comparison of NADH, NADPH, and FAD binding pockets in CpFNR from *C. pasteurianum*. Panel 1 illustrates the residues involved in the docking of NADH, colored in salmon, including residues 315, 320, 311, 86, 312, 85, and 84. Panel 2 highlights NADPH binding residues in red, specifically residues 316, 195, 319, 317, 313, and 86. Panel 3 compares the binding residues between NADH and NADPH. Panel 4 indicates the FAD binding residues, highlighted in yellow, including residues 310, 312, and 86. Panel 5 compares the relative binding locations of NADH, NADPH and FAD, shedding light on the potential structural basis for differences in binding affinity and catalytic function.

### The CpFd–CpFNR interface is governed by electrostatic complementarity

ClusPro docking of CpFd onto CpFNR identified an interface composed of 14 residues on CpFNR and 12 residues on CpFd (Fig. 6). The interface is stabilised by a combination of electrostatic interactions (Asp6 on CpFd with Arg188 on CpFNR; Glu55 on CpFd with His5 on CpFNR), hydrogen bonds and hydrophobic contacts. This pattern of acidic ferredoxin residues pairing with basic FNR residues is consistent with the electrostatic complementarity observed in the *Anabaena* FNR–ferredoxin complex [7,8,10,23–26]. The comprehensive spatial analysis of all binding partners (Fig. 7) placed the FAD binding region centrally between the NAD(P)H and CpFd binding domains, consistent with the established electron transfer relay.

**Figure 6.**
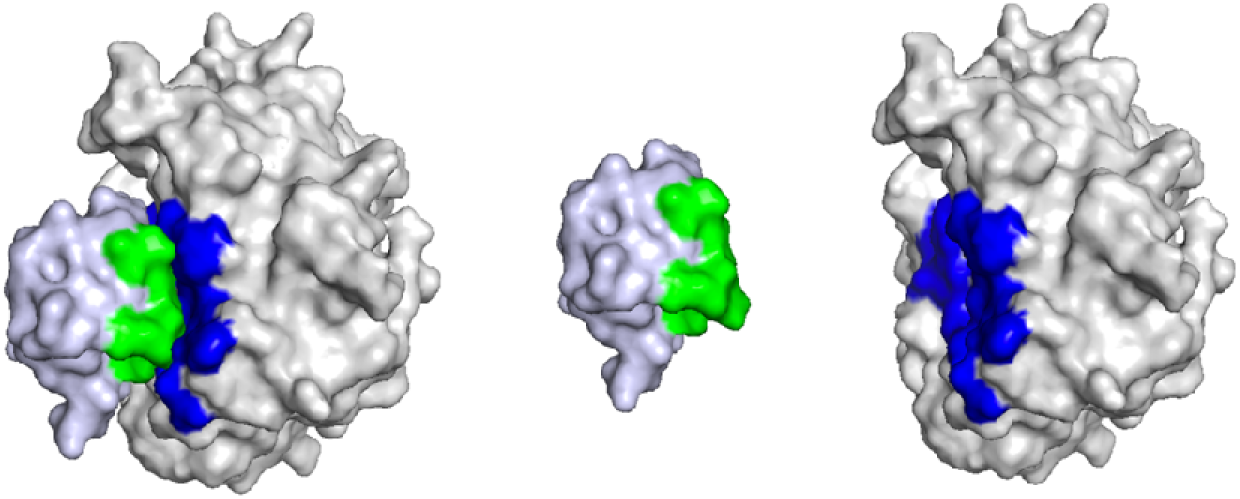
Docking regions of CpFd (light blue) and CpFNR (grey). The CpFNR interface is highlighted in blue, while the CpFd interface is shown in green.

**Figure 7.**
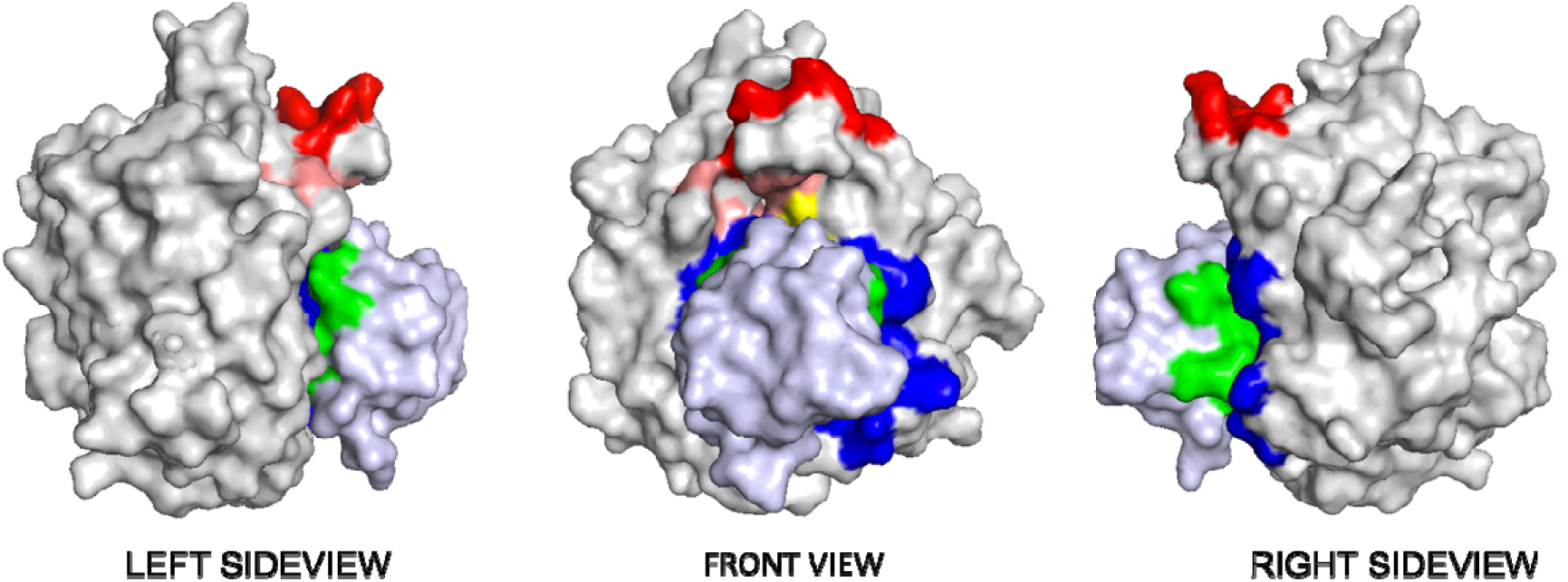
The relative locations of NADH (salmon), NADPH (red), and FAD (yellow) binding pockets on CpFNR (grey) and the CpFd docking domains (light blue). The CpFNR interface is highlighted in blue, while the CpFd interface is shown in green.

### Structural rationalisation of beneficial mutations through loop flexibility

Mapping of mutation sites onto the CpFNR structure (Fig. 8) revealed that S202A and T185A are located at opposite ends of a random coil spanning residues 186–199 that connects the NAD(P)H and CpFd binding regions. The S202A mutation disrupts hydrogen bonds with Asn199, potentially increasing the conformational flexibility of this coil and propagating effects to the secondary structures formed between Ala309 and Cys324, where critical NAD(P)H interactions occur. The T185A mutation, positioned at the opposite end of the same coil, disrupts hydrogen bonds with His215 and Thr7, similarly destabilising the coil. Thus, although S202A and T185A are located at opposite ends of the 186–199 coil, both mutations converge on the same mechanistic outcome: increased coil flexibility leading to enhanced substrate affinity.

**Figure 8.**
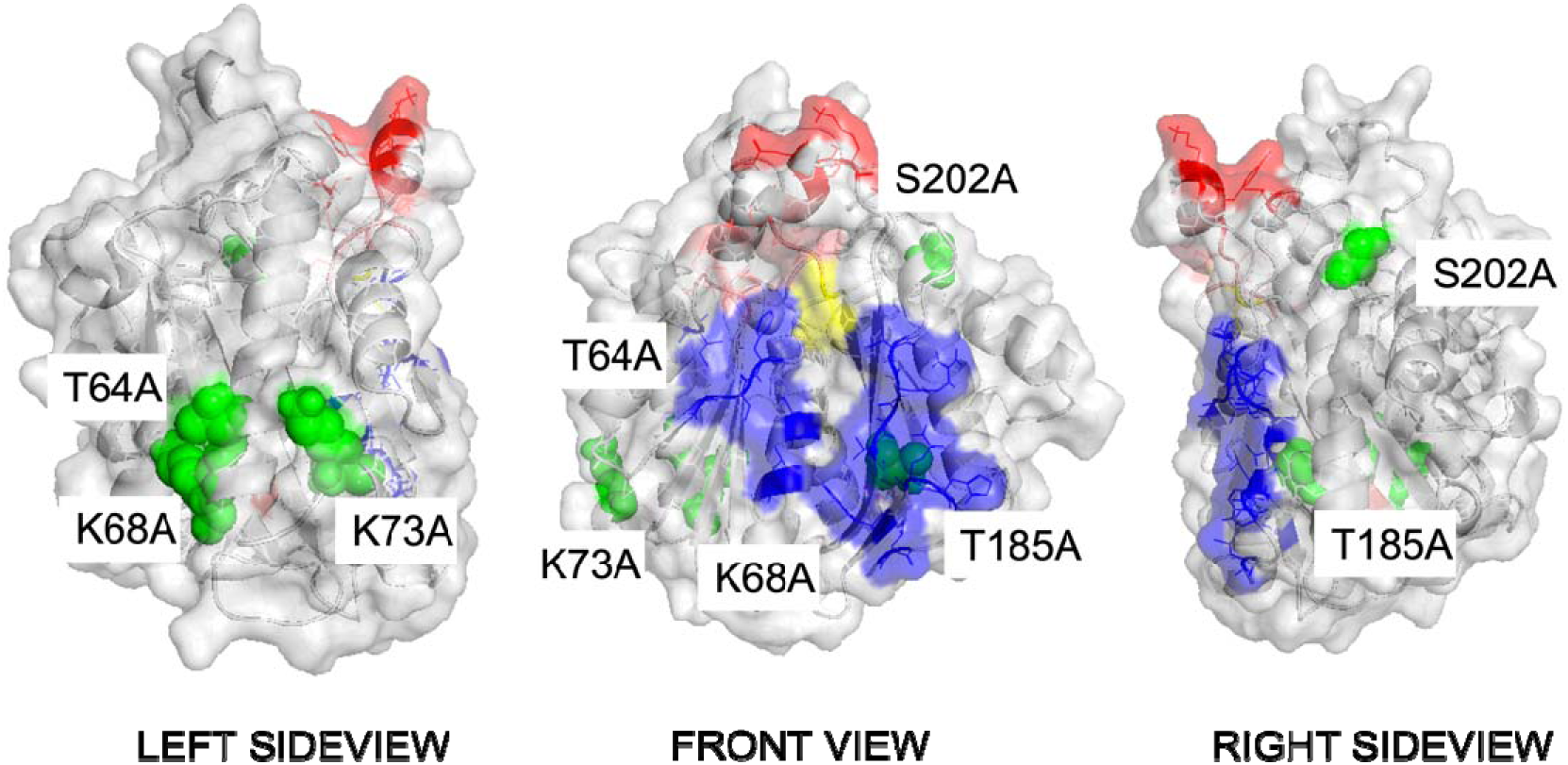
Locations of alanine scanning mutations on CpFNR relative to cofactor and ferredoxin binding domains. Mutation sites are shown as spheres.

The T64A mutation operates through a distinct mechanism. Located within an α-helix (residues Cys50–Gly69), Thr64 forms two hydrogen bonds with Glu60. Loss of the side-chain hydroxyl upon mutation to alanine may distort the helix, altering the angle of the downstream structure and directly affecting NAD(P)H binding residues Leu84 and Met85, while also propagating effects through β-sheet elements to CpFd binding residues.

## Discussion

In the present study, we have identified the gene encoding ferredoxin–NADP^+^ reductase activity in *C. pasteurianum*, resolving a gap that has persisted since the activity was first reported by Jungermann and co-workers in 1973 [12]. The finding that this activity resides in AQ984_05830, a protein annotated as a sporulation factor, was unexpected and has several implications for our understanding of both the FNR superfamily and the developmental biology of spore-forming clostridia.

The exceptionally low *k*_cat_ values of CpFNR (0.007–0.02 min^−1^) compared to canonical FNRs (49–349 s^−1^) [3] raise the question of whether this enzyme serves a metabolic or regulatory function. It has been proposed that optimal redox conditions promote the physiological processes necessary for endospore formation in clostridia, including chromosome segregation and assembly of the protective spore coat [35,36]. In this context, the low catalytic rate of CpFNR would be consistent with a role in fine-tuning the intracellular NAD(P)H/NAD(P)^+^ ratio during sporulation rather than sustaining bulk NADPH production. This hypothesis is reminiscent of the role played by redox-sensing regulators such as Rex in other clostridia, which modulate gene expression in response to changes in the NADH/NAD^+^ ratio. Future studies examining the expression profile of *AQ984_05830* during the sporulation programme and the phenotype of deletion mutants will be required to test this model.

The genome-wide ferredoxin survey (Fig. 3) offers a broader context for the discovery of CpFNR. The presence of numerous ferredoxin-like proteins, rather than a single carrier, is consistent with a metabolic architecture in which electrons liberated from molecular hydrogen are not funnelled to one destination but are instead distributed across several ferredoxins, each feeding a distinct physiological process. In *C. pasteurianum*, H oxidation by the [FeFe]-hydrogenase generates a pool of reduced ferredoxin that must be apportioned among competing demands—nitrogen fixation, proton reduction and hydrogen recycling, biosynthetic reductions, and redox homeostasis. The identification of CpFNR adds a previously missing branch to this scheme: a route by which reduced ferredoxin can regenerate NADPH. Viewed this way, the multiplicity of ferredoxins is not redundancy but specialisation, allowing the organism to route a shared supply of hydrogen-derived reducing equivalents toward whichever pathway is required at a given metabolic or developmental stage. The discovery of a sporulation-associated protein moonlighting as an FNR fits naturally within such a flexible, ferredoxin-centred electron-distribution network.

From a structural perspective, several features distinguish CpFNR from canonical FNRs. The overlap between FAD and NAD(P)H binding regions on the second loop (residues 309–313) contrasts with the spatially separated binding domains observed in well-characterised FNRs [4,5,18–20], and the **<u>G</u>**ALL**<u>S</u>**PL**<u>S</u>**motif on this loop differs from the GXSXXS motif identified in FdR9 [4]. These structural differences, combined with the sporulation annotation, raise the possibility that CpFNR represents a distinct branch of the FNR superfamily that has been co-opted for a developmental function. It would be of considerable interest to determine whether homologues of AQ984_05830 with FNR activity exist in other spore-forming clostridia, which could illuminate the evolutionary origins of this moonlighting function.

The identification of beneficial mutations (T64A, T185A, S202A) that improve catalytic efficiency through increased flexibility of the 186–199 coil region is consistent with recent work demonstrating the importance of loop dynamics in enzyme catalysis [29–31]. The convergence of S202A and T185A on the same mechanistic outcome—despite their location at opposite ends of the coil—suggests that this structural element functions as a conformational switch that modulates access to the NAD(P)H binding site. Engineering of this coil region, guided by the structural insights presented here, may provide a route to improving the catalytic efficiency of CpFNR for biotechnological applications in cofactor regeneration systems [28].

## Conclusions

In summary, we have identified a novel ferredoxin–NADP^+^ reductase from the *C. pasteurianum* genome through systematic bioinformatic screening. The enzyme, AQ984_05830, is annotated as a sporulation protein and exhibits FNR activity with a *k*_cat_ orders of magnitude lower than canonical FNRs. Alanine scanning mutagenesis and structural analysis revealed distinct mechanisms by which mutations enhance catalytic efficiency, converging on increased flexibility of a coil region connecting the cofactor and ferredoxin binding domains. The discovery that a sporulation-associated protein moonlights as an FNR suggests an unrecognised role for redox regulation during endospore formation and expands the known functional diversity of the FNR superfamily. More broadly, the coexistence of CpFNR with an array of ferredoxin carriers points to a hydrogen-driven electron-distribution network in which multiple ferredoxins channel reducing equivalents toward specialised metabolic and developmental functions.

## Acknowledgements

The authors acknowledge funding from the Department of Energy to James Swartz (SDACJ-1-1257268).

## Author contributions

WW conceived the study, performed experiments and wrote the manuscript. QL contributed to discussion and editing. JRS conceived the study, supervised the work and edited the manuscript.

## Data availability statement

All data supporting the findings of this study are available within the article and its supplementary materials.

